# Mice carrying paternal knockout of imprinted *Grb10* do not show compulsive behaviour

**DOI:** 10.1101/2020.04.03.016014

**Authors:** Kira DA Rienecker, Alexander T Chavasse, Kim Moorwood, Andrew Ward, Trevor Humby, Anthony R Isles

**Affiliations:** MRC Centre for Neuropsychiatric Genetics and Genomics, Neuroscience and Mental Health Research Institute, School of Medicine, Cardiff University, Hadyn Ellis Building, Maindy Road, Cardiff, UK; Department of Biology and Biochemistry, University of Bath, Building 4 South, Bath, BA2 7AY, UK; School of Psychology, Cardiff University, Tower Building, Park Place, Cardiff, UK

**Author notes:** Corresponding author *ARI.

## Abstract

Mice lacking paternal expression of imprinted *Grb10* show a number of social behaviour deficits, including an enhanced allogrooming phenotype. However, this could also index compulsive behaviour, and the increased whisker barbering seen in *Grb10*^+/p^ mice has been suggested to be indicative of a trichotillomania-type behaviour. Here we test whether compulsive behaviour is a more general phenotype in *Grb10*^+/p^ mice by examining marble burying. We also examined the mice for potentially confounding anxiety phenotypes using the elevated plus maze (EPM). *Grb10*^+/p^ mice showed no difference from wild-type littermate controls on any measure in the marble burying test at any age. However, *Grb10*^+/p^ mice were more willing to enter, spend time on, and head-dip on the open arm of the EPM than wild-type mice. These data suggest that *Grb10*^+/p^ mice are not generally more compulsive, and that the enhanced allogrooming is probably indicative of altered social behaviour. Furthermore, the altered behaviours seen on the EPM adds to other published findings suggesting that *Grb10* has a role in mediating risk-taking.

## INTRODUCTION

Imprinted genes, defined by their parent-of-origin specific epigenetic marks and monoallelic expression, are highly expressed in the brain and linked to behaviour. Paternal expression of imprinted *Grb10* is prominent in monoaminergic regions of the midbrain and influences a range of behaviours (Dent et al., 2020; Dent et al., 2018; Garfield et al., 2011). In particular, *Grb10* paternal knockout mice (*Grb10*^*+/p*^) mice have an enhanced allogrooming phenotype (Garfield et al., 2011) which has been inferred to indicate social dominance, but could also index compulsive behaviour (Haig and Ubeda, 2011). Altered grooming behaviours are often used to model compulsivity in mice, as features such as a focused affected area have compelling similarities to trichotillomania (compulsive hair plucking). Genetic knockout models of compulsivity have linked facial over-grooming and anxiety phenotypes to monoaminergic dysregulation in the cortex and striatum (Wood et al., 2018). Likewise, monoaminergic neurotransmitter systems where *Grb10* is expressed are implicated in the pathophysiology of obsessive compulsive disorder (OCD), an anxiety disorder characterized by compulsive behaviour and obsessive thinking (Albelda & Joel, 2012). Nevertheless, compulsivity has not been explicitly assayed in *Grb10*^*+/p*^ mice.

Here, we set out to specifically assay general compulsive behaviour in the *Grb10*^*+/p*^ mouse. The marble burying test (MBT) has good face validity for repetitive and compulsive behaviour, and detects differences between treatment conditions known to manipulate relevant neurotransmitter systems (Albelda and Joel, 2012). To dissociate from known confounding effects of anxiety in the MBT, we also examined behaviour on the Elevated Plus Maze (EPM). Our previous observations of *Grb10*^*+/p*^ colonies indicates that barbering behaviour emerges with age (Rienecker et al., 2020). Therefore, we employed a cross-sectional design to assess any potential compulsivity phenotype at two ages (6 and 10 months). Overall, we found no evidence to support a compulsivity in the *Grb10*^*+/p*^ mice, but data from EPM measures supports previous findings indicating that mice lacking expression of paternal *Grb10* are more willing to take risks.

## MATERIALS AND METHODS

*Grb10* heterozygous knockout mice on a B6CBAF1/J background were as previously described (Rienecker et al., 2020). The study used two separate groups of *Grb10*^*+/p*^ and wild-type (WT) littermate mice (6 and 10 months old at the start of testing, Supplementary Fig. 1) in a cross-sectional design. Mice progressed through the marble burying test (MBT), followed by the elevated plus maze (EPM). All procedures were conducted in accordance with the requirements of the UK Animals (Scientific Procedures) Act 1986, under the remit of Home office license number 30/3375 with ethical approval at Cardiff University.

In the Marble Burying Task (MBT) animals were tested in an arena with a deep sawdust layer where one half contained eight red marbles. Mice were placed in the arena and allowed to freely explore the arena for 30 minutes. Behaviours were scored by Ethovision, apart from the number of marbles displaced, half buried, and buried, which were recorded manually every 5 minutes.

The Elevated Plus Maze (EPM) was used to assess anxiety. The maze consisted of two bisecting white Perspex arms (43cm x 8 cm, l x w) at right angles to each other, fixed to a stand 45 cm high. Opposing pairs of arms were designated “Closed arms” (with walls) and “Open arms” (without walls). Animals were placed in a closed arm and allowed to freely explore the maze for 5 minutes. Behaviour was scored by the Ethovision detection system, apart from head dips, which were scored manually.

All statistical analyses were carried out using SPSS 26.0 for windows (IMP Corp., USA). Ethovision measures in the MBT and EPM were analysed using ANCOVA, with between-subjects independent variable GENOTYPE and SEX, co-varied for AGE (6 and 10 months). Manually scored behaviours from the MBT (marbles displaced, half-buried, and totally buried) were analysed by repeated measures ANCOVA with an additional within-subject factor TIME (5 minute intervals for 30 minutes). Benjamini-Hochberg False discovery rate (FDR) was applied to correct for multiple testing (i.e. measures and analyses within a given test).

## RESULTS

### *Grb10*^+/p^ mice show no differences in marble burying

*Grb10*^*+/p*^ and wild-type litter mate controls showed no difference in the cumulative rate of marbles being buried in the MBT, even before correction for multiple testing (ANOVA, main effect of GENOTYPE F_1, 171_=0.22, p=0.641, partial η^2^=0.001; Figure 1A). Although there was a difference between males and females (ANOVA, main effect of SEX F_1, 171_=5.86, p=0.017, partial η^2^ = 0.001; Figure 1A), this did not survive correction for multiple testing and there was no interaction with GENOTYPE (F_1, 171_=2.31, p = 0.13, partial η^2^ = 0.013).

**Figure 1.**
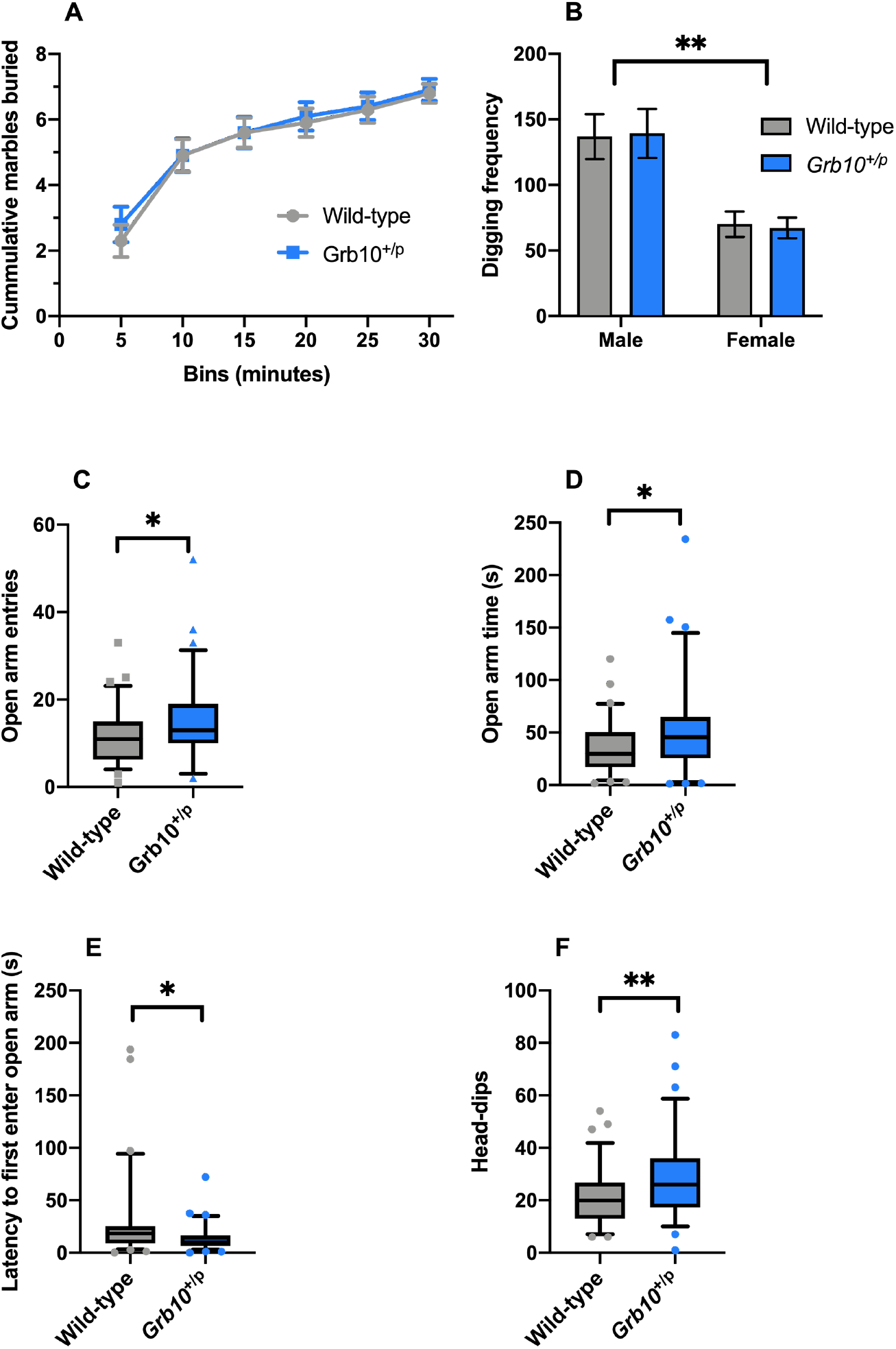
Behaviour of *Grb10*^+/p^ in the MBT and EPM. Cumulative number of marbles buried (**A**) and digging frequency (**B**) does not differ between *Grb10*^+/p^ and wild-type mice. However, there was a significant difference between males and females in digging frequency that survived FDR correction. On the EPM, *Grb10*^+/p^ mice made more entries into (**C**) and spent more time on (**D**) the open arm than wild-types. *Grb10*^+/p^ were also significantly quicker to first enter the open arm (**E**) and made more head-dips (**F**) that wild-type mice. Data report estimated marginal mean with 5^th^ and 95^th^ percentile, apart from **A** and **B** that show estimated marginal mean ±SEM.

This main measure is supported data by ‘digging frequency’ that also suggest no difference between *Grb10*^*+/p*^ and wild-type mice (ANOVA, main effect of GENOTYPE F_1, 171_=0.03, p=0.873, partial η^2^=0.0002; Figure 1B) and no interaction between GENOTYPE and SEX (F_1, 171_=0.26, p=0.612, partial η^2^=0.002; Figure 1B). However, there was a significant effect of SEX (F_1, 171_=107.6, p=0.0001, partial η^2^=0.38; Figure 1B) that survived FDR (p=0.004).

### *Grb10*^+/p^ mice are not more anxious on the EPM, but show ancillary phenotypes

The EPM was included to differentiate between anxious and compulsive behaviour in the MBT. Although no differences were seen in the MBT, *Grb10*^*+/p*^ mice showed a number of behavioural differences on the EPM. Specifically, they made more entries into the open arm (ANOVA, main effect of GENOTYPE, F_1,151_=6.40, p=0.012, partial η^2^ = 0.042; Figure 1C); spent more time on the open arm (ANOVA, main effect of GENOTYPE, F_1,151_=7.43, p=0.007, partial η^2^=0.048; Figure 1D); were quicker to enter the open arm (ANOVA, main effect of GENOTYPE, F_1,151_=9.98, p=0.002, partial η^2^=0.064; Figure 1E; and made more head-dips over the edge of the open arm (ANOVA, main effect of GENOTYPE, F_1,151_=10.80, p=0.001, partial η^2^ = 0.068; Figure 1F) than wild-type littermates. All these test survived FDR correction (p=0.019; p=0.015; p=0.013; p=0.010, respectively). There were no differences between males and females, and no interactions between GENOTYPE and SEX that survived FDR correction for any measure (see Supplementary materials).

## DISCUSSION

We have previously reported that *Grb10*^*+/p*^ mice have altered social stability behaviour (Garfield et al., 2011; Rienecker et al., 2019), including enhanced allogrooming, or whisker barbering (Garfield et al., 2011). While allogrooming correlates with other measures of social dominance (Wang et al., 2011), it is also used to model compulsivity in rodents (Albelda and Joel, 2012) and it has been suggested that trichotillomania and obsessive-compulsive-like behaviour may explain this phenotype in *Grb10*^*+/p*^ mice (Haig and Ubeda, 2011). Therefore, we systematically assessed compulsivity in *Grb10*^*+/p*^ mice using the marble burying test (MBT) and included the elevated plus maze (EPM) to assay potentially confounding differences in anxiety. *Grb10*^*+/p*^ mice showed no evidence of abnormal compulsive behaviours in the MBT, discounting this explanation of altered allogrooming behaviour. However, behaviour on the EPM was altered, with *Grb10*^*+/p*^ mice being more willing to enter, spending more time on, and making more head-dips on the open arm than wild-type litter mates. Taken together with previous findings (Dent et al., 2020; Dent et al., 2018), we suggest these data provide further support for the idea that *Grb10*^*+/p*^ mice are more willing to take risks.

We found no difference in marble burying measures in male and female *Grb10*^+/p^ mice. The lack of evidence for compulsive-type behaviour in the MBT is supported by our previous findings in the Stop-Signal Reaction Time (SSRT), task where *Grb10*^*+/p*^ mice show no differences in the SSRT compared to controls (Dent et al., 2018). The SSRT is generally related to ‘stopping impulsivity’, as the task measures the capacity to inhibit an already initiated response, but a tendency towards simple stereotyped movements, as in compulsive behaviour, can impair performance on this test (Robbins et al., 2012). Collectively these data, coupled with the finding that *Grb10*^*+/p*^ and wild-type mice regrow barbered whiskers following social isolation (Garfield et al., 2011; Rienecker et al., 2019), indicate that the enhanced barbering associated with *Grb10*^*+/p*^ mice results not from a general compulsive behavioural phenotype, but is probably indicative of altered social behaviour.

We included the EPM in our experiment as the MBT alone cannot differentiate between compulsive and anxious behaviours (Albelda and Joel, 2012), and to expand our anxiety assessments in this cross-sectional study by including both sexes and two age points. On the EPM, *Grb10*^+/p^ mice spent more time on the open arm and displayed an increased willingness to enter the open arm as indicated by a higher total number of entries and a reduced latency to first enter. These findings would suggest a decreased anxiety phenotype, and so are at odds with those reported previously for the open field and light-dark box tests where no difference was seen between *Grb10*^+/p^ and wild-type mice (Garfield et al., 2011). However, behaviour on the EPM, in particular the increase in head-dipping, may also be interpreted as measure of risk-taking behaviour (Toledo-Rodriguez and Sandi, 2011).

The data presented here, together with previous studies (Dent et al., 2018), indicate *Grb10*^*+/p*^ mice do not have a generalised compulsive behavioural phenotype. Thus, altered allogrooming is not confounded by compulsivity behaviour, and instead supports findings of altered social stability in *Grb10*^*+/p*^ mice (Rienecker et al., 2020). Additionally, data from the EPM suggests *Grb10*^*+/p*^ mice are more willing to explore the open arm. Coupled with findings from the delayed reinforcement (Dent et al., 2018) and Predator Odour Risk-Taking tasks (Dent et al., 2020), we suggest these EPM data are another indication that paternal *Grb10* normally acts to reduce risk-taking behaviour.

## Data Availability

All data is freely available at: https://osf.io/n9kmz/

