## Supplementary materials for "Mice carrying paternal knockout of imprinted *Grb10* do not show compulsive behaviour"

### DETAILED MATERIALS AND METHODS

#### *Subjects*

*Grb10* heterozygous knockout mice on a B6CBAF1/J background were created using a LacZ:neomycin gene-trap cassette interrupting exon 7 as previously described (Garfield et al., 2011). The colony was derived via embryo transfer from a colony in Bath and breeding stock was maintained with either a B6CBA F1/crl line from Charles River (C57BL/6J:CBA/CaCrl F1 mice, the first generation progeny of a cross between female C57BL/6J and male CBA/CaCrl mice) or with an in house mixed B6CBA F1/crl x B6CBA F1/J background. Experimental animals (F2) were generated by crossing heterozygous $Grb10^{+/-}$ males with wildtype females in order to generate litters of wild-type and $Grb10^{+/p}$ pups. Overall, the study used three separate groups of male wild-type and $Grb10^{+/p}$ mice (2, 6, and 10 months old at the start of testing, Supplementary Fig. 1) in a cross-sectional design. However, here we focus only on the 6- and 10-month age groups, as these are the ages at which whisker barbering appears (Rienecker et al., 2020). Mice progressed through social hierarchy tests (Rienecker et al., 2020) before the marble burying test (MBT), followed by the elevated plus maze (EPM).

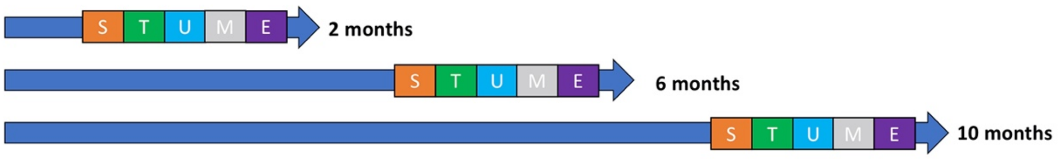


**Supplementary Figure 1. Schematic outlining the order of testing.** Three cohorts, aged 2, 6, and 10 months were tested for stranger-encounter tube test (S), social-encounter tube test (T), urine marking test (U; males only), marble burying test (M), and EPM (E). Testing ended at the nominal age of the cohort and was confined to a 4 week period to prevent overlap between cohorts. Only marble burying and EPM are reported in this paper. Results from the remaining tests were reported previously (Rienecker et al., 2020) where a version of this figure appears.

Animals were weaned and housed as previously described (Rienecker et al., 2020). All groups of mice were housed in environmentally enriched cages (cardboard tubes, shred-mats, chew sticks) of 1-5 adult mice per cage. Cages were kept in a temperature and humidity-controlled vivarium (21 ± 2˚C and 50 ± 10% respectively) with a 12-hour light-dark cycle (lights on at 7:00 hours, lights off at 19:00 hours). All mice had *ad libitum* access to standard rodent laboratory chow and water. Cages were cleaned and changed once a week at a regular time and day of the week for minimal disruption. All procedures were conducted in accordance with the requirements of the UK Animals (Scientific Procedures) Act 1986, under the remit of Home office license number 30/3375 with ethical approval at Cardiff University.

#### *Behavioural Procedures*

All experiments were carried out in a quiet room with one overhead light (15 lux), and behaviour was analysed using Ethovision video-tracking software (V3.0.15, Noldus Information Technology, Netherlands), via a camera placed centrally over each piece of apparatus. We used a set of quantitative descriptors about the movement and location of subjects, determined by the location of the greater body-proportion of subjects (12 frames/s). We calibrated tracking using non-experimental mice of the same body size and coat colour as experimental subjects. For each experiment, we designated appropriate virtual zones related to the apparatus and tracked parameters such as distance moved, time spent/zone, zone entries, velocity, and latency to enter a zone. One cage of four mice was carried into the testing room at a time and remained until all cage mates had individually completed the task. Between cages in the marble burying task, 1/3 of the sawdust was removed and replaced with fresh material. We cleaned marbles and the EPM apparatus with 70% alcohol solution between each trial. Mice were handled as little as possible up until one week prior to the start of behavioural testing; then the researcher who would perform the behavioural tests handled the mice daily for 5 days, recording weight and barbering status.

#### *Marble Burying Task (MBT)*

The MBT was conducted using previously published methods (Doe et al., 2009). Mice were placed in an arena (40 x 24 x 11, l x w x h in cm) three-quarters filled with levelled sawdust and covered by a Perspex lid with narrow gaps on either 24 cm end of the box. Eight red marbles (10mm diameter) were placed in a set pattern of three rows (2 x 3 marbles and a centre row of 2 marbles) in one half of the arena (“Marbles Zone”). To begin the trial, mice were placed in the “Start Zone” and allowed to freely explore the arena. Mice were recorded in the arena for 30 minutes with number of marbles displaced, half buried, and buried recorded manually every 5 minutes, and an overall total for the session was determined. Digging and grooming times were manually scored throughout the trial. Following the trial, the sawdust was turned over and fresh marbles were placed in the “Marbles Zone”. The main measures of the MBT were “marbles buried”, “marbles half-buried”, “marbles displaced”, “velocity”, “total time digging”, “percent time in ‘start’ zone”, “percent time in “marbles’ zone”, “transitions”, “total time digging”, and “total time grooming”. “Marbles buried” and “Marbles half-buried” were combined to provide a cumulative overall score.

#### *Elevated Plus Maze (EPM)*

The EPM was conducted to control for confounding effects of anxiety on the marble burying test. The maze consisted of two bisecting white arms (43cm x 8 cm, l x w) at right angles to each other, was made of white Perspex, and was fixed to a stand 45 cm high. Opposing pairs of arms were designated “Closed arms” (with 17cm high walls) and “Open arms” (without walls). All arms opened onto a centre square of 8 cm x 8 cm, designated the “Middle Zone”. To begin the 5-minute trial, a mouse was placed in a closed arm and allowed to freely explore. The main measures reported were “entries to the open arm”, and “time in open arms”, “latency to first enter open arm” and “head-dips”. The Ethovision detection system recorded movement, while head dips over the edge were scored manually.

**Statistics**

Benjamini-Hochberg FDR was calculated by running the following Syntax in SPSS 26.0

DATA LIST free / p (F5.3).
BEGIN DATA
**.152 .093 .055 .035 .044 .017 .001** (*P-values of interest*)
END DATA.
SORT CASES by p (a).
COMPUTE i=$casenum.
SORT CASES by i (d).
COMPUTE q=.05.
COMPUTE m=max(i,lag(m)).
COMPUTE crit=q*i/m.
COMPUTE test=(p le crit).
COMPUTE test=max(test,lag(test)).
FORMATS i m test(f8.0) q (f8.2) crit(f8.6).
VALUE LABELS test 1 'Significant' 0 'Not Significant'.
LIST

**RESULTS**

| **Measure** | **Factor** | **F-value** | **P-value** | **Partial $\eta^2$** |
|---|---|---|---|---|
| Cumulative marbles buried | Time | 8.813 | **3.62E-8** | 0.049 |
| | Genotype | 0.218 | 0.641 | 0.001 |
| | Sex | 5.857 | 0.017 | 0.033 |
| | Genotype*Sex | 2.314 | 0.130 | 0.013 |
| Digging frequency | Genotype | 0.025 | 0.874 | 0.0002 |
| | Sex | 107.01 | **8.77E-20** | 0.385 |
| | Genotype*Sex | 0.257 | 0.613 | 0.002 |

**Supplementary Table 1** Statistics for measures in the marble burying test. Emboldened P-values are those that survived Benjamini-Hochberg FDR correction for multiple testing.

| **Measure** | **Factor** | **F-value** | **P-value** | **Partial $\eta^2$** |
|---|---|---|---|---|
| Open arm entries | Genotype | 6.402 | **0.012** | 0.042 |
| | Sex | 0.019 | 0.890 | 0.0001 |
| | Genotype*Sex | 2.083 | 0.151 | 0.014 |
| Time on open arm | Genotype | 7.430 | **0.007** | 0.048 |
| | Sex | 0.123 | 0.726 | 0.001 |
| | Genotype*Sex | 0.769 | 0.382 | 0.005 |
| Latency to first enter open arm | Genotype | 9.979 | **0.002** | 0.064 |
| | Sex | 1.266 | 0.262 | 0.009 |
| | Genotype*Sex | 0.204 | 0.652 | 0.001 |
| Head-dips | Genotype | 10.799 | **0.001** | 0.068 |
| | Sex | 1.630 | 0.204 | 0.011 |
| | Genotype*Sex | 1.137 | 0.288 | 0.008 |

**Supplementary Table 2** Statistics for measures in the elevated plus maze. Emboldened P-values are those that survived Benjamini-Hochberg FDR correction for multiple testing

**Data Availability** All data is freely available at: https://osf.io/n9kmz/

**Funding** This work was funded by Wellcome grant 105218/Z/14/Z. ARI is part of the MRC Centre for Neuropsychiatric Genetics and Genomics (G0801418).

**Competing interests** The authors declare they have no competing interests.
